# Cytomegalovirus pentamer dependent endocytic cellular infection utilizes redundant entry receptors in the guinea pig model

**DOI:** 10.64898/2026.05.07.723586

**Authors:** Yushu Qin, K. Yeon Choi, Alistair McGregor

**Affiliations:** Department of Microbial Pathogenesis & Immunology, Texas A&M University, Health Science Center, College of Medicine, Bryan, Texas, United States of America

## Abstract

2.

The guinea pig with guinea pig cytomegalovirus (GPCMV) is the only small animal model for congenital CMV (cCMV). GPCMV cell entry is dictated by specific viral gH/gL-based complexes: gH/gL/gO trimer (direct entry); pentamer complex, PC (endocytic entry). GPCMV gB as the fusogenic protein is also essential for all entry pathways. PDGFRA and NRP2 are receptors for direct and endocytic virus entry respectively based on strain 13 animal fibroblast ATCC cell line studies. All non-fibroblast guinea pig cell lines are derived from Dunkin-Hartley animals, the focus of cCMV studies. GPCMV infection of Dunkin-Hartley embryo fibroblasts (GEFh) and epithelial cells were compared. Knockout of PDGFRA on GEFh cells prevented GPCMV(PC-) direct entry but not endocytic GPCMV(PC+) infection, demonstrating both pathways of infection. Fibroblast generated virus poorly infected epithelial cells compared to epithelial virus stock, which exhibited full tropism to all cell types. Guinea pig epithelial cell lines are NRP2-positive and PDGFRA-negative requiring PC for GPCMV infection. Epithelial and GEFh cells, but not strain 13 fibroblasts, additionally expressed ThBD. In immunoprecipitation assays, PC and ThBD interacted unlike CD46 receptor candidate targeting gH/gL. Double-knockout of NRP2/ThBD in epithelial cells impaired infection unlike single knockouts. Individual ectopic species-specific receptor expression restored infection on double-knockout epithelial (NRP2/ThBD) and fibroblast (PDGFRA/NRP2) cell lines. Knockout of NRP2/ThBD receptors did not enhance GPCMV neutralization by gB antibodies on PDGFRA-negative cells demonstrating a limitation of a gB vaccine strategy. Overall, GPCMV and HCMV similarity for receptors and cell tropism maintains the translational importance of this model.

**Impact Statement:** A CMV vaccine is a high priority as congenital cytomegalovirus (cCMV) is a leading cause of hearing loss and cognitive impairment in newborns. CMV species-specificity requires animal studies to utilize species-specific virus. The guinea pig is the only small animal model for cCMV and guinea pig cytomegalovirus (GPCMV) encodes functional HCMV homolog glycoprotein complexes for cell entry including a gH/gL-based PC for endocytic cell entry. The viral fusogenic gB glycoprotein is required for GPCMV infection of all cell types but antibody based gB vaccines fail to fully protect against endocytic infection. The PC is potentially an important vaccine antibody target, but PC-based cell entry is only partially understood and poorly characterized for GPCMV. Identifying GPCMV cell entry receptors is critical to the understanding of virus tropism and disease especially since the PC is necessary for cCMV. Correlation with HCMV improves translational impact of guinea pig based cCMV intervention strategies.

## 5. Introduction

Human cytomegalovirus (HCMV) is a ubiquitous pathogen, and congenital infection leads to symptomatic disease in newborns including cognitive impairment and sensorineural hearing loss (SNHL) (1–3). Furthermore, congenital CMV (cCMV) can be a risk factor for miscarriage or stillbirths, fetal intrauterine growth retardation, preterm birth and autism (4, 5). Globally, cCMV occurs in approximately 1-5% of live births and natural convalescent immunity to the virus does not prevent infection by a new strain with the potential for non-primary cCMV (6). In the US, long-term health care related to cCMV amounts to billions of dollars and a vaccine is considered a high priority (7). HCMV species-specificity complicates preclinical animal model studies, which requires the use of species-specific animal CMV. The guinea pig is the only small animal model for cCMV with guinea pig cytomegalovirus (GPCMV) causing disease in newborn pups (8–11). Various cCMV intervention strategies have been evaluated in this model with varying levels of success (11, 12).

Viral glycoprotein complexes on the virion are neutralizing antibody targets and are of interest as CMV vaccine antigens. Specific heterologous gH/gL-based glycoprotein complexes interact with cell receptors to enable viral cell entry by one of two pathways: [1] direct; or [2] endocytic (13). Direct cell entry is dependent upon viral complexes gB, gM/gN, and gH/gL/gO (14–19). An additional HCMV gH/gL-based pentamer complex (PC) in association with gB enables endocytic infection of epithelial, endothelial and myeloid cells (20, 21). Guinea pig CMV (GPCMV) encodes essential homolog glycoprotein complexes for direct cell entry (gB, gH/gL/gO, gM/gN) and these are neutralizing target antigens (11, 22–24). GPCMV encodes a homolog PC (gH/gL/GP129/GP131/GP133) necessary for endocytic pathway of infection (23). Cellular platelet-derived growth factor receptor alpha (PDGFRA), present mainly on fibroblasts, has been identified as receptor for HCMV direct cell entry (25). Similarly, guinea pig PDGFRA enables GPCMV direct infection via interaction with viral gH/gL/gO trimer but viral cell entry also required gB (24, 26). Various guinea pig cell lines have recently been established where GPCMV epithelial and endothelial infection requires PC for endocytic entry as they lack PDGFRA (24, 27, 28). However, PDGFRA can act as a universal receptor for GPCMV direct cell entry in all cell types by ectopic expression (27, 28).

The gB glycoprotein is an immunodominant antibody target and essential for infection (22, 23, 29–34). Therefore, gB is considered a key component of any vaccine against cCMV. However, a gB subunit vaccine, despite evoking high antibody titers, attains approximately 50% efficacy in preclinical or clinical trials (11, 35–37). In both HCMV and GPCMV, gB antibodies are less effective at neutralizing infection of non-fibroblast cells (33, 38–40). Other antigens are being explored as vaccine candidates including the PC (21, 41, 42). The process of PC-dependent cell entry is only partially understood with several human receptor candidates identified for HCMV including: neuropilin-2 (NRP2), OR14I1, CD147, CD46, and thrombomodulin (ThBD) (43–48). GPCMV PC is necessary for infection of cells lacking PDGFRA including epithelial, endothelial, and trophoblast cells as well as macrophage (11, 23, 27, 28, 49, 50). Consequently, the PC is highly important for GPCMV dissemination and congenital infection (23, 27, 28, 49). Indeed, PC antibodies in a vaccine strategy against GPCMV improves neutralizing-antibody titers and contributes to enhancement of protection against cCMV (24, 51). Given the importance of GPCMV PC to virus infection and the development of an effective CMV vaccine or therapeutic, it was critical to gain a better understanding of viral endocytic infection by identifying GPCMV candidate PC cellular receptors.

Previously, guinea pig PDGFRA KO cells remained susceptible to GPCMV(PC+) infection via the endocytic pathway (24, 26, 52). Analysis of available epithelial, endothelial and fibroblast guinea pig cell lines identified several potential GPCMV PC entry receptors (NRP1, NRP2 and CD147) universally expressed in all cell lines (52). However, only guinea pig NRP2 interacted with viral PC (52). Double-knockout of PDGFRA and NRP2 completely blocked GPCMV fibroblast (GPL) infection demonstrating the importance of these receptors for GPCMV infection pathways (52). However, there were limitations associated with these initial studies with a focus on fibroblast cells. Additionally, the GPL fibroblast ATCC cell line used in the studies was of strain 13 guinea pig origin and not Dunkin-Hartley strain animals. Importantly, the majority of GPCMV pathogenicity, vaccine and cCMV studies are carried out in Hartley guinea pigs and all established non-fibroblast cell lines are derived from Hartley strain animals. Consequently, it was important to establish a Hartley-derived fibroblast cell line for comparative studies. Additionally, virus receptor research was extended to GPCMV epithelial cellular infection as this cell type is important for viral shedding (saliva or urine) (23, 53), as well cCMV infection via placental trophoblasts and amniotic sac epithelial cells (27, 49) with GPCMV infection requiring PC. We hypothesized that targeted PC receptor knockout on epithelial cells would impair or block GPCMV endocytic infection. Candidate guinea pig receptors were evaluated by: [1] interaction of receptor with GPCMV gH/gL, PC or viral trimer; [2] impact on GPCMV infection by single or double cellular receptor knockout; [3] ectopic expression of receptors for GPCMV infection on knockout cell lines. Results demonstrated that GPCMV utilizes a similar strategy to HCMV for PC entry receptors in a species-specific manner. However, GPCMV can only maximally utilize the PC endocytic pathway if virus stocks are generated on non-fibroblast cell lines (eg. epithelial cells) as GPCMV passed once on fibroblast cells lacked full tropism to epithelial cells. Additionally, results demonstrate a limitation of gB vaccine antibodies in epithelial cell GPCMV neutralization despite the essential nature of gB and studies with PC receptor knockout cell lines. Overall, studies illustrate the similarity of HCMV and GPCMV infection increasing the translational relevance of CMV intervention strategies evaluated in the guinea pig model.

## 6. Materials & Methods

### 6.1 Cells, viruses, oligonucleotides and genes

GPCMV (strain 22122, ATCC VR682 or clinical strain TAMYC) were propagated on various guinea pig cell lines previously established and characterized (23, 27, 28, 49). Guinea pig cell lines used in the study include guinea pig (strain 13) fibroblast lung cells (GPL; ATCC CCL 158) and various guinea pig (Dunkin-Hartley) epithelial cells including: renal epithelial (REPI); amniotic sac (GPASE); guinea pig umbilical cord endothelial cells (GPUVEC); trophoblasts (TEPI) (23, 27, 28, 49). Additionally, an embryotic guinea pig (Dunkin-Hartley) fibroblast cell line (GEFh) was established following previously described protocol (54). GPCMV BAC derived virus (PC+/PC-/GFP+/-) were generate as previously described (23). GFP-tagged wild type HSV-1 (17+ strain) was previously described (55). Recombinant defective adenovirus (Ad5) vectors (E1 and E3 deleted) were previously described encoding individual components of the PC (gH, gL, GP129, GP131, GP133) and additional recombinant Ad vector encoded gO (33, 56). All ORFs were under HCMV IE enhancer control with 3’ SV40 polyA sequence. High titer CsCl gradient purified recombinant defective adenovirus virus stocks (10^12^ TDU/ml) were generated by Welgen Inc.

(MA). All oligonucleotides were synthesized by Sigma-Genosys (The Woodlands, TX). Synthetic codon optimized genes were developed for candidate guinea pig receptors: NRP2; ThBD; CD46; OR14I1 (Genscript). Guinea pig ORFs were derived from guinea pig NCBI genome sequence (*Cavia porcellus* annotation release 103 GCF_000151735.1): NRP2 (XM_013157537); ThBD XP_005008421.2; and CD46 NP_001166399.2. ORFs additionally incorporated a C-terminal FLAG epitope tag. The ORFs were cloned into pcDNA3.1(+) vector (Invitrogen) under HCMV IE enhancer promoter control to enable transient expression in plasmid transfected cells with protein expression verified by Western blot analysis. Human NRP2, ThBD and PDGFRA pcDNA3.1(+) based expression vectors were commercially validated expression constructs (Genscript).

### 6.2 Ethical Approval Statement

Cell lines used in the described studies were previously established and characterized as described in previous publications (23, 27, 28, 49). Prior approach for the acquisition of guinea pig (Dunkin-Hartley) tissues to establish cell lines was carried out under IACUC (Texas A&M University) permit. Guinea pig fibroblast lung cell line (GPL; ATCC CCL-158) and a colorectal carcinoma epithelial cell line (GPC-16; ATCC CCL-242) were obtained from ATCC. A guinea pig embryo fibroblast cell line (GEFh) was generated from a pregnant (Dunkin-Hartley strain) animal mid-gestation following a previously described protocol (54). All prior study animal procedures were carried out in strict accordance with the recommendations in the “Guide for the Care and Use of Laboratory Animals of the National Institutes of Health.” Animals were observed daily by trained animal care staff, and animals that required care were referred to the attending veterinarian for immediate care or euthanasia. Terminal euthanasia was carried out by lethal CO_2_ overdose followed by cervical dislocation in accordance with IACUC protocol (Texas A&M University) and NIH guidelines. Generated Dunkin-Hartley guinea pig cell lines from the McGregor lab used in the study include: embryo fibroblast lung cells (GEFh): renal epithelial cells (REPI); amniotic sac (GPASE); umbilical cord endothelial cells (GPUVEC); aorta endothelial; and trophoblasts (TEPI). Establishment of guinea pig cell lines fully described in earlier publications (23, 27, 28, 49, 54).

### 6.3 Western blot assays

Western blots were carried out on cell lysates separated by 4-20% SDS-PAGE under denaturing conditions and performed as previously described (22, 23, 33). For western blots, anti-epitope tag primary antibodies were used at 1/1000: FLAG (Novus Biological); GFP (Santa Cruz Biotechnology); His (Bethyl): MYC-c (Novus Biologicals) (23, 24). Secondary antibodies: anti-mouse or anti-rabbit IgG HRP-linked secondary antibodies for western blot (Cell Signaling) were used at 1/2000 (24). FLAG epitope was used to detect CD147, NRP2, GP133 and gO protein expression in transduced/transfected cell monolayers. Additionally, species cross-reactive antibodies were used for the detection of NRP2 (R&D Systems), CD147 (Antibodies Online), ThBD (Antibodies Online) and PDGFRA (R&D Systems).

### 6.4 Immunoprecipitation assays

Immunoprecipitation (IP) assays were carried out on plasmid transfected or recombinant Ad transduced guinea pig cells using commercial GFP-trap (ChromoTek) following manufacturer’s protocols with inclusion of protease inhibitor cocktail (Pierce) in cell lysates. Samples were subsequently analyzed by SDS-PAGE (4–20% gradient gel) and western blot using specific anti-epitope tag antibodies: FLAG (Novus Biological); GFP (Santa Cruz Biotechnology); His (Bethyl): MYC-c (Novus Biologicals); and CD147, NRP2 and ThBD specific antibodies. Appropriate secondary anti-mouse or anti-rabbit HRP conjugate (Cell Signaling Technology) were also used following standard western blot protocol as previously described (23).

### 6.5 Bafilomycin A1 inhibition of pH-dependent endocytosis on PDGFRA KO cells

Bafilomycin A1 (Baf A1) inhibition of GPCMV infection was carried out following a previously described protocol (52). Fibroblast cells (wild type or PDGFRA KO) in six-well plates were pretreated with complete media containing either 0 or 100 nM Baf A1 (Sigma) for 1 h at 37 °C, followed by GPCMV(PC+) GFP-tagged virus infection (MOI=0.5 p.f.u. cell−1) for 1 h at 37 °C. All further incubations were performed at the same concentration as pretreatment in complete media. Cells were fixed at 24 h post-infection with 4% PFA (15 min) and subsequently overlaid with PBS and stored at 4 °C as sealed plates. Infected cells were identified by GFP fluorescent microscopy. Infection was set up in triplicate. Counts were made of GFP-positive cells in random fields. Statistical analysis was performed using Student’s t-test on the percent of cells infected in 30 random fields of view, each containing ∼100 cell nuclei, for each condition. The number of treated cells infected was represented as a percentage of the number of infected untreated cells.

### 6.6 Virus neutralization assays

GPCMV neutralization assays (NA_50_) were performed on GPL fibroblasts, PDGFRA KO fibroblasts, renal epithelial and receptor KO epithelial cells (REPI NRP2 KO and REPI ThBD/NRP2 DKO) with GPCMV(PC+) virus stocks generated on renal epithelial cells. Neutralization assays were performed with historical pooled sera from a specific group of trimeric capable AdgB vaccinated animals (33). Serially diluted sera were incubated with approximately 1 × 10^5^ pfu GPCMV(PC+) in media containing 1% rabbit complement (Equitech Bio) for 90 min at 37 °C before infecting GPL KO, REPI or REPI KO cells for 1 hour. For neutralization on GPL cells, 1 × 10^3^ pfu GPCMV(PC+) was used. Infected cells and supernatant were collected on day 4 then titrated on GPLs. Final neutralizing antibody titer was the inverse of the highest dilution producing 50% or greater reduction in plaques compared to virus only control. NA_50_ were performed from each sample three times concurrently with the same virus stocks between groups.

### 6.7 CRISPR/Cas9 mutagenesis knockout strategy

#### Guinea pig PDGFRA gene knockout of fibroblast GEFh cells

Knockout of PDGFRA in GEFh cells was carried out as previously described (24). The sequence of the guinea pig PDGFRA (gpPDGFRA) gene (NCBI gene accession number 100726209) was based on the guinea pig NCBI genome sequence (Cavia porcellus annotation release 103 GCF_000151735.1), and predicted introns and exons were additionally identified via use of the genome with Ensembl accession number ENSCPOG00000011782. Exon 2 of gpPDGFRA was targeted for mutagenesis by use of the CRISPR/Cas9 strategy. Exon 2-specific guide RNA (gRNA) was designed via an online program (www.rgenome.net) to avoid off-target sites. DNA sequences were cloned under US6 promoter control in three separate gRNA expression plasmids (pCas-Guide-EF1a-GFP; OriGene): pR1 (5′-GGTGTGGGCCGCCGAGGCGT-3′), pR2 (5′-TCTGGGAGAGTTCCCCGACG-3′), and pR3 (5′-CGTTTCTGATGTCCACGTCG-3′). GEFh cells in 6-well plates were transduced with defective lentivirus expressing Cas9 under HCMVIE enhancer control (Origene) to enable expression of Cas9 and subsequently transfected with the gRNA expression plasmids (pR1, pR2, and pR3). At 2 days post-transfection, the cells were reseeded in 96-well plates by limiting dilution to generate individual cell lines. PDGFRA gene knockout cell lines were screened by exon 2 PCR sequencing of extracted genomic DNA. Genomic extraction was carried out with a DNeasy extraction kit (Qiagen), and PCR was performed using primers Fpd (5′- CTGAGCCTAATCTGCTGCCAGCTTTCG-3′) and Rpd (5′- CGGCACGGTAGATGTAGATATGC-3′). Western blotting of total cell lysate for specific modified cell lines verified the loss of gpPDGFRA expression using primary antibody Goat anti-mouse PDGFRA antibody (R&D Systems). The PCR products of the wild-type GEFh and PDGFRA mutant knockout cell lines were cloned into the TA cloning vector (Invitrogen) and sequenced as previously described (28). Alignment of the PDGFRA exon 2 DNA sequence with wild-type and KO cells was previously shown (24) and modification to PDGFRA in exon 2 of knockout Dunkin-Hartley embryo fibroblasts cell line was as previously described (24). PDGFRA knockout GEFh cells were designated GEFh PDGFRA KO.

#### Generation of single (NRP2 or ThBD) and double knockout renal epithelial (REPI) cell line (ThBDNRP2 DKO)

Single or double-gene knockout renal epithelial cell lines were generated following the same protocol described above for knockout of PDGFRA with Cas9 positive cell transfected with appropriate sets of gRNA expression plasmids. Cells were seeded by limiting dilution and cell lines were subsequently screened by PCR exon sequencing for DNA modification and western blot for loss of target protein expression to confirm single or double-gene knockout using specific primers for NRP2 or ThBD.

#### NRP2 and ThBD gRNA

Guinea pig NRP2 (XM_013157537) and ThBD (XP_005008421.2) gene sequence was derived from guinea pig NCBI genome sequence (*Cavia porcellus* annotation release 103 GCF_000151735.1) as described for PDGFRA (52). Predicted introns and exons were additionally identified via use of the genome analysis with Ensembl. Exon specific gRNAs were designed via online program www.rgenome.net/be-designer/ to avoid off-target sites. DNA sequences were cloned under US6 promoter control in three separate gRNA expression plasmids (pCas-Guide, OriGene) for each target gene. NRP2 gRNA plasmids: nR1. 5’ATGGACCATCGAATCTCCG3’; nR2. 5’GATGATCTCCATCTTGGGTT3’; nR3.

5’AGGGTCATGCTCCAGGTCAA3’; nR4. 5’ATGGGGACAGCGAGTCCG3’.

ThBD gRNA plamsids : nT1 ATGCTCGGGGTGCTGCTCCT: nT2 CCTGGGGCTCCTGGTGCCCG; nT3 TCCTGGTGCCCGCGGACCCG

Gene knockouts confirmed by comparative Western blots of cell lysates for receptor protein expression and exon 2 PCR sequencing of knockout locus. Confirmed single KO cell line designated NRP2 KO or ThBD KO. Double-knockout cell lines were designated as ThBD/NRP2 DKO REPI cells for knockout of both NRP2 and ThBD.

### 6.8 BLAST protein alignment

Alignment and predicted identity for candidate receptors (NRP2, ThBd, and CD46) between guinea pig and human counterparts was carried out with predicted amino acid sequence of guinea pig and human proteins based from NCBI data base with alignment carried out with NCBI BLASTp program (https://blast.ncbi.nlm.nih.gov/Blast.cgi). NCBI accession numbers for predicted proteins: Guinea pig NRP2 (XP_013012991); ThBD XP_005008421.2; CD46 NP_001166399.2; Human NRP2 (NP_957718); ThBd NP_000352.1; CD46 NP_758868.1.

### 6.9 Statistical analysis

All statistical analyses were conducted with GraphPad Prism (version 7) software. Student’s t test (unpaired) with significance taken as a *p* value of <0.05 or as specified in the figure legends. Analysis of Variance (ANOVA) one-way was performed to compare multiple groups in a study.

## 7. Results

### 7.1 Hartley strain fibroblasts express multiple GPCMV entry receptors including ThBD

Original GPCMV (22122) research utilized Hartley guinea pig embryo fibroblast cells for evaluation of virus growth in tissue culture (57). However, studies from the mid-1990s onwards utilized an established ATCC embryo fibroblast cell line (GPL ATCC CCL 158) for virus growth and titration experiments because of an ability of cells to be contact inhibited as a confluent monolayer and these cells easily supports virus, BAC and plasmid transfections (11). However, this cell line was derived from strain 13 animals. In contrast, all current non-fibroblast epithelial and endothelial established cell lines are derived from Hartley guinea pigs, which is the animal strain used in all cCMV and virus pathogenicity studies. Consequently, a Hartley embryo fibroblast cell line (designated GEFh) was established to enable contrasting studies with Hartley non-fibroblast cell lines.

Western blot analysis of GEFh cell lysate demonstrated that this new cell line expressed both PDGFRA and NRP2 receptors with similar levels of expression to GPL cells (Fig 1A). However, unlike GPL cells, GEFh fibroblasts also expressed an additional candidate receptor, ThBD (Fig 1A). A previously described CRISPR Cas9 knockout strategy was utilized to ablate PDGFRA expression in GEFh cells and establish a novel PDFRA knockout cell line (GEFh PDGFRA KO). The modified cell line was verified by Western blot analysis of cell lysate for loss of PDGFRA protein expression (Fig 1A) and genetic modification of the cell line by PDGFRA exon 2 by sequencing (data not shown). One-step growth curve studies of GPCMV(PC+) and GPCMV(PC-) virus on GEFh and GEFh PDGFRA KO cells demonstrated that GPCMV(PC+) virus had similar growth kinetics on both cell lines (Fig 1B). In contrast, GPCMV(PC-) virus had normal growth kinetics on GEFh cells but was unable to infect GEFh PDFRA KO cells (Fig 1C). Ectopic expression of PDGFRA on GEFh PDGRA KO cells restored GPCMV(PC-) infection to normal kinetics (Fig 1C). Unlike wild type GEFh cells, GPCMV(PC+) infection of GEFh PDGFRA KO cells were dependent upon an acidic pH flux for endocytic cell entry as Bafilomycin A1 was capable of blocking infection (data not shown) as previously observed for PDGFRA KO GPL cells (52), renal and trophoblast epithelial cells (23, 49). A control infection of HSV-1 on GEFh and KO cell line produced similar levels of virus at 2 dpi on both cell lines (Fig 1D). This demonstrated that specific impact upon infection related to GPCMV(PC-) and loss of direct entry receptor PDGFRA. Results were similar to that obtained for GPL PDGFRA knockout fibroblasts (52). The limited ability of GEFh PDGFRA KO cells to support GPCMV(PC-) infection implied that ThBD likely functioned as a PC receptor similar to HCMV (47). A BLAST comparison of guinea pig and human ThBD demonstrated 63% identity to human counterpart. In addition to GEFh cells expressing ThBD, Western blot analysis of Hartley epithelial and endothelial cell lysates demonstrated ThBD protein expression on established cell lines (Fig 2A). However, ThBD protein levels were higher in endothelial cells compared to epithelial cells (Fig 2A). ThBD is a type-1 integral membrane protein acting as an anticoagulant cofactor for thrombin in humans and guinea pigs (58). Consequently, given the function of ThBD it was perhaps not unexpected to find higher levels of ThBD expression on endothelial cells. In order to determine if ThBD interacted with GPCMV glycoprotein trimer (gH/gL/gO) or PC viral glycoprotein complexes, separate immunoprecipitation (IP) assays were carried out. IP assays utilized a GFP-trap strategy directed to GFP tagged gH and transient expression of trimer or PC and FLAG epitope tagged plasmid expressed ThBD as previously described (23, 52). Results demonstrated a specific interaction only with PC and no interaction with viral trimer (Fig 2C & D). It was concluded that ThBD is likely a PC receptor for GPCMV.

**Figure 1.**
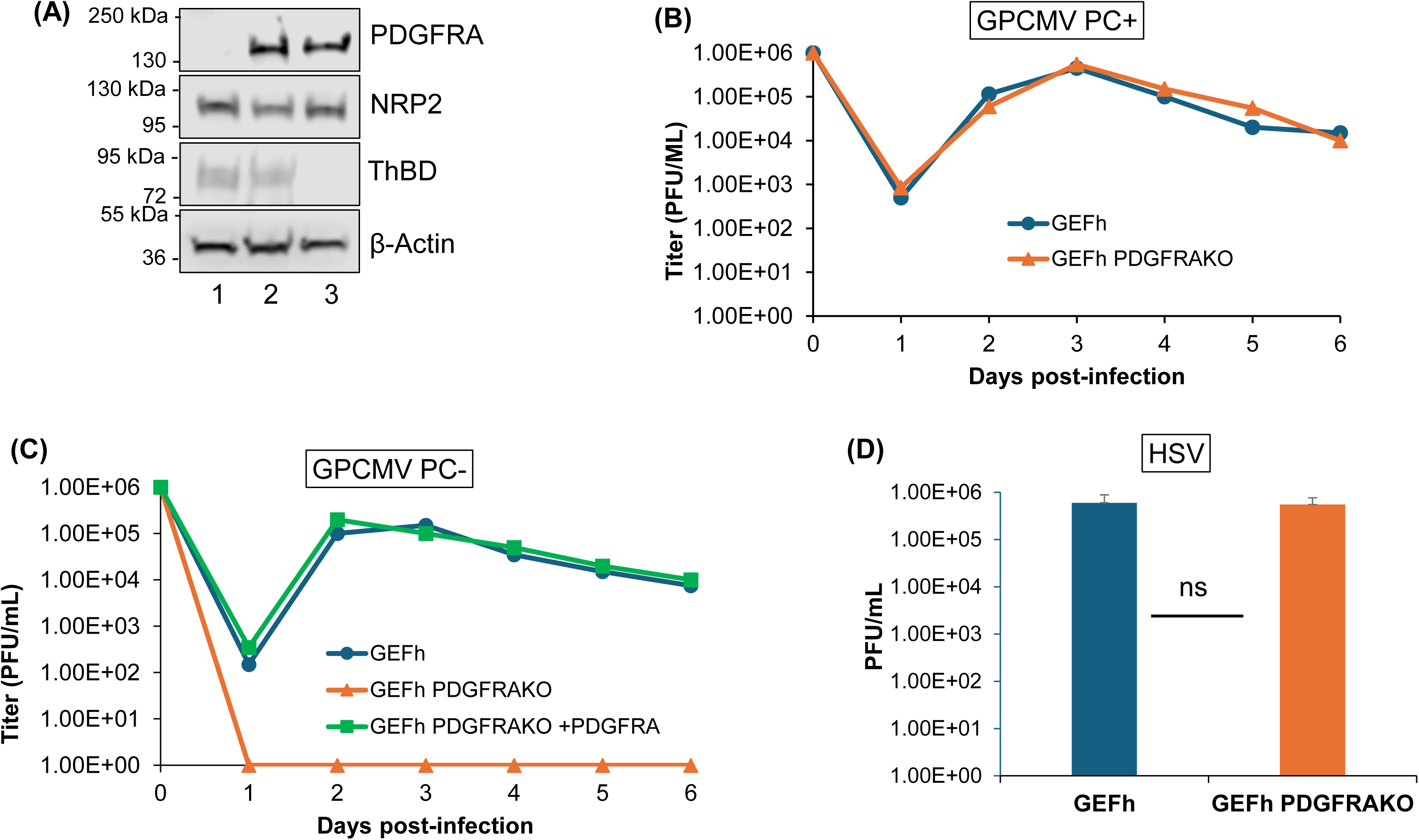
Characterization of Dunkin-Hartley derived embryo fibroblast cells (GEFh). (A) Western blot analysis of cell lysate for expression of PDGFRA, NRP2 and ThBD cellular receptors: GEFh PDGFRA-KO (lane 1) GEFh (lane2); and GPL (lane 3). (B) One-step growth curve of GPCMV (PC+) wild type virus (MOI 1 pfu/cell) on GEFh (blue circle) vs GEFh PDGFRA-KO (orange triangle). (C) Growth curve of GPCMV (PC-) virus on GEFh (blue circle) vs GEFh PDGFRA-KO (orange triangle) and GEFh PDGFRA-KO rescue transfected with PDGFRA expression plasmid (green square). (D) HSV-1 infection on GEFh (blue) vs GEFh PDGFRA-KO (orange). MOI = 1 pfu/cell, harvested 2 DPI. Statistical analysis Student *t* test, ns = non-significant.

**Figure 2.**
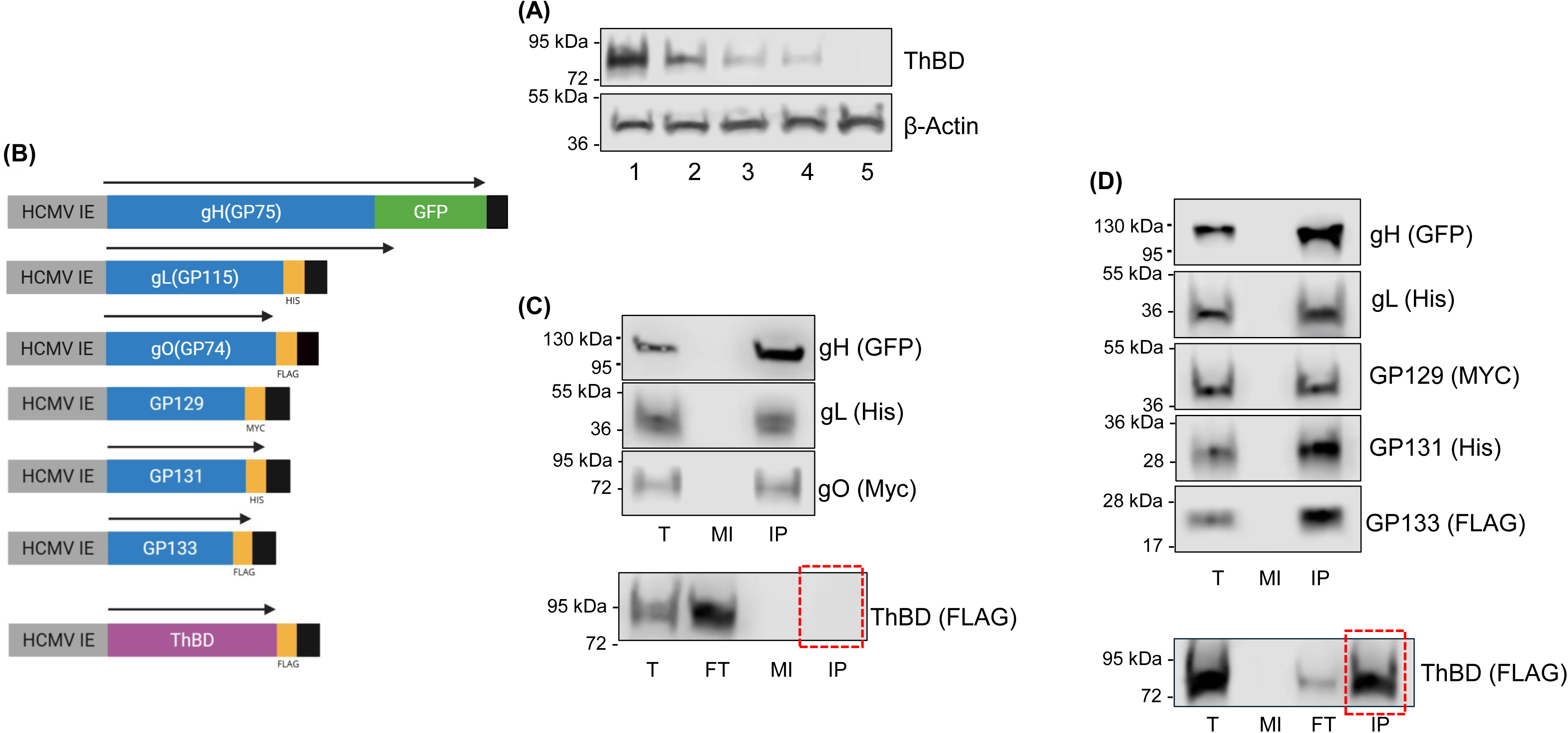
Evaluation of ThBD receptor expression in guinea pig cells and interaction of GPCMV gH based trimer or PC with ThBD via immunoprecipitation (IP) assay. (A) ThBD expression in guinea pig cell lines. Western blot analysis of cell lysate for expression of ThBD: Endothelial GPUVEC (lane1); Aorta endothelial (lane 2); REPI epithelial (lane 3); TEPI trophoblast (lane 4); GPL (lane 5). (B) Diagram of specific expression plasmids used for individual trimer and PC components and ThBD. (C)-(D) Determination of (C) gH based trimer or (D) PC interaction with plasmid transfected ThBD(FLAG) co-expression followed by GFP-trap IP as previously described (25). Primary antibodies used in Western blot analysis: α-GFP (gH); α-His (gL), α-FLAG (gO), α-MYC (GP129); α-His (GP131); α-FLAG (GP133); α-FLAG (ThBD). Samples analyzed as total cell lysate or processed for IP assay followed by Western blot analysis. T: total cell lysate. F: flowthrough. MI: mock control lysate. IP: immunoprecipitated product.

### 7.2 Epithelial cell NRP2 and ThBD expression and impact on GPCMV infection

Recently, we demonstrated that NRP2 was a PC receptor for GPCMV and that double-knockout of PDGFRA and NRP2 receptors on GPL cells rendered the double-knockout cell line resistant to GPCMV infection by blocking both direct and endocytic entry pathways (52). This result demonstrated the importance of NRP2 receptor for endocytic entry. Additionally, double-knockout of PDGFRA and CD147 dramatically impaired GPL infection with CD147 having an indirect effect on endocytic cell entry (52). A limitation of prior studies was that they were performed on GPL fibroblast cells. Additionally, GPL cells only expressed NRP2 and it was not possible to determine if endogenous NRP2 or ThBD was more important for GPCMV endocytic cell entry. Consequently, receptor entry knockout studies were expanded to epithelial cells. We hypothesized that since epithelial cell infection was fully dependent upon PC for infection via endocytic pathway, a knockout of both identified GPCMV PC receptors (NRP2 and ThBD) would likely block or highly impair epithelial cell GPCMV infection. ThBD is expressed at roughly similar levels on all Hartley epithelial cell lines including REPI, TEPI and amniotic sac (Figure 3A). NRP2 receptor was expressed at similar levels on guinea pig epithelial cells (52). A CRISPR-based knockout strategy was pursued on renal epithelial (REPI) cells to knockout individual receptors (NRP2 or ThBD) or to generate a double-knockout receptor (NRP2 and ThBD) mutant REPI cell line. CRISPR Cas9 based knockout strategy was carried out as described in materials and methods (52) targeting exon 2 of each gene respectively. Isolated knockout cell lines were verified for loss of expression of receptor protein by Western blot of cell lysate (Fig 3B & C) as well as sequencing of knockout locus on exon 2 of NRP2 or ThBD gene (data not shown). Single NRP2 knockout REPI cells and double-knockout NRP2/ThBD cells were designated NRP2 KO and ThBD/NRP2 DKO respectively. Single-knockout ThBD cells were designated ThBD KO. Comparative Western blot analysis of cell lysates from wild type REPI cells or knockout cell lines demonstrated the loss of NRP2 expression in NRP2 KO and ThBD/NRP2 DKO cell lines (Fig 3B). Loss of ThBD expression was similarly confirmed by Western blot analysis (Fig 3C). Beta-actin was used as a lane load control for Western blot analysis of cell lysates (Fig 3A-C).

**Figure 3.**
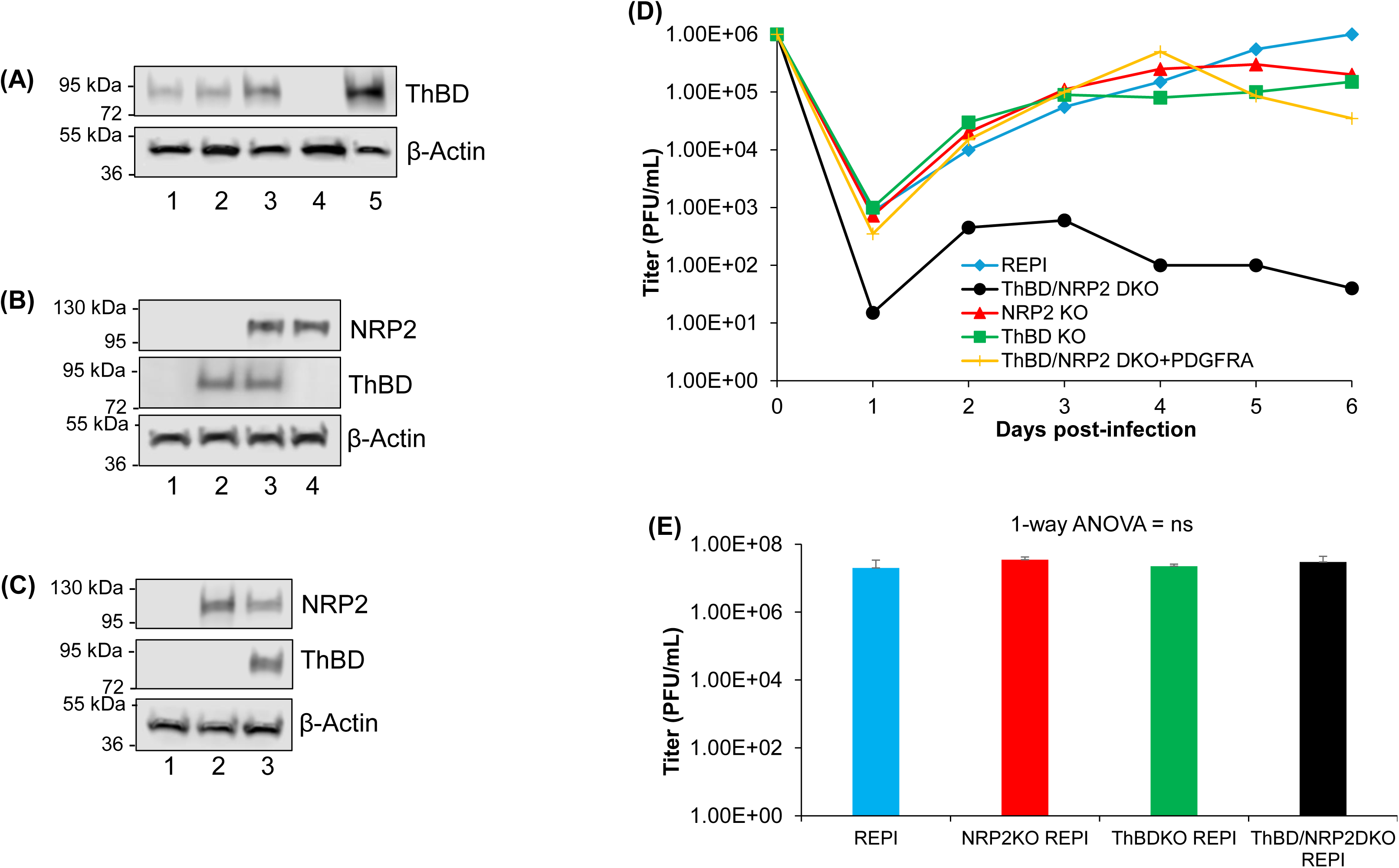
Characterization of NRP2 and ThBD single and double-knockout guinea pig epithelial REPI cell lines. (A) Western blot analysis on various epithelial cell lines for ThBD expression. Lanes: 1. GPASE; 2. TEPI; 3. REPI; 4. GPL; 5. HEK293. Gel loading confirmed by β-actin expression. (B) Western blot analysis of NRP2 KO and ThBD/NRP2 DKO cells for expression of NRP2 or ThBD. Lanes: 1. ThBD/NRP2 DKO REPI; 2. NRP2KO REPI; 3. REPI; 4. GPL. Gel loading confirmed by β-actin expression. (C) Western blot analysis of ThBD KO and ThBD/NRP2 DKO cells for expression of NRP2 or ThBD. Lanes: 1. ThBD/NRP2 DKO REPI; 2. ThBD KO REPI; 3. REPI. Gel loading confirmed by β-actin expression. (D) Growth curve of GPCMV (PC+) wildtype virus on REPI (blue diamond); NRP2 KO (red triangle); ThBD KO (green square); ThBD/NRP2 DKO (black circle); ThDB/NRP2 DKO + PDGFRA (yellow slash). (E) HSV-1 infection on REPI (blue); NRP2 KO REPI (red); ThBD KO REPI (green); ThBD/NRP2 DKO REPI (black). MOI = 1, harvested 2 DPI. Statistical analysis one-way ANOVA, ns = non-significant.

After confirming knockout of receptors on the various REPI cell lines, GPCMV(PC+) infection experiments were performed on confluent monolayers on six well plates to evaluate infection by one-step growth studies (MOI 1 pfu/cell). Results demonstrated that all cell monolayers could be infected by GPCMV (Figure 3D). Single-knockout of ThBD or NRP2 had no real impact on viral infection. In contrast, double-knockout of ThBD and NRP2 resulted in impaired infection on DKO cells by several log values compared to single-knockout or wild type REPI cells (Fig 3D). In contrast to PDGFRA/NRP2 double-knockout on GPL cells, which resulted in cells fully resistant to GPCMV infection. Control HSV-1 infection produced similar levels of infection on single and double-knockout cells compared to wild type REPI cells (Fig 3E). Results demonstrated that loss of two key entry receptors had a specific impairment only on GPCMV infection. Potentially REPI cells express an additional receptor to enable GPCMV infection despite NRP2/ThBD DKO. Prior studies demonstrated that all guinea pig cell lines expressed CD147 and that this receptor had an indirect effect on GPCMV PC infection (52). Potentially additional knockout of CD147 on ThBD/NRP2 DKO REPI cells could further reduce cell susceptibility to infection, but this awaits additional study.

Individual ectopic expression of guinea pig ThBD or NRP2 on ThBD/NRP2 DKO cells restored GPCMV infection to levels equivalent to that of wild type REPI cells (Fig 4A-C) confirming the impairment to infection was associated with receptor loss and that only one receptor was necessary to restore endocytic infection levels. In contrast, ectopic expression of human receptor counterparts on DKO REPI cells did not rescue infection (Fig 4D-F) and indicated a basis for species-specific barrier related to GPCMV infection. An established guinea pig epithelial adenocarcinoma ATCC cell line (GPC16) was additionally evaluated for virus infection. Although this cell line expressed both NRP2 and ThBD (Fig S1A), a comparative one-step growth curve of GPCMV(PC+) on REPI and GPC16 cells demonstrated that the adenocarcinoma cell line was highly resistant to GPCMV infection compared to REPI cells (Fig S1B) and plasmid expression of receptors (PDGFRA, NRP2 or ThBD) did not enhance GPCMV infection (data not shown). In contrast, HSV-1 infected both REPI and GPC16 cells with slightly higher progeny virus obtained on GPC16 cells (Fig S1C). Results demonstrate, the GPC16 cell line is supportive of HSV-1 but not GPCMV despite expression of NRP2 and ThBD. Consequentially, it is assumed that additional factors can impair CMV infection with tumor derived cell lines being restrictive to GPCMV as similarly observed for HCMV.

**Figure 4.**
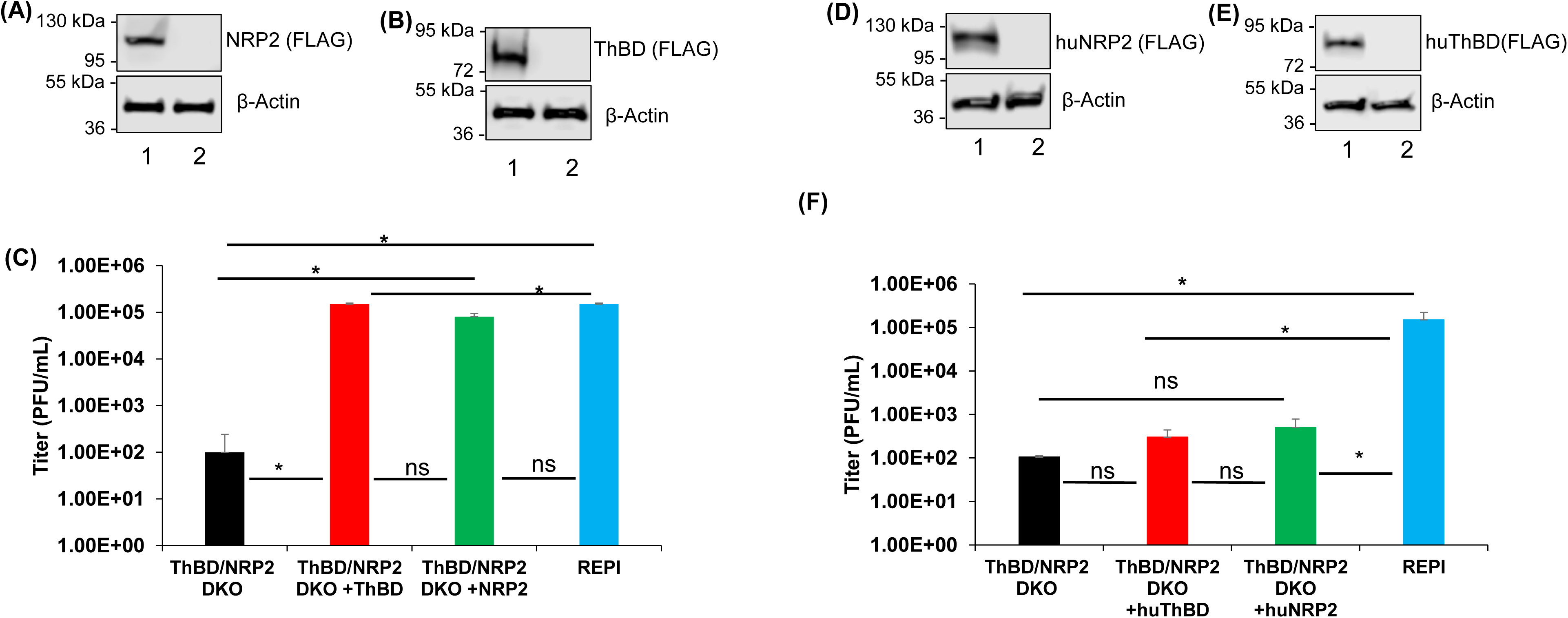
Complementation of GPCMV infection on guinea pig epithelial cell receptor knockout cells by ectopic expression of human or guinea pig NRP2 or ThBD. (A)-(C) Guinea pig receptors and GPCMV infection on ThBD/NRP2 DKO REPI cells. (A) Western blot analysis of REPI cells transfected with plasmid NRP2(FLAG) expression vector. Upper blot, NRP2 detected with α-FLAG primary antibody. Lanes: 1, NRP2(FLAG) plasmid-transfected cells; 2, control REPI cells. Lower blot, β-Actin Western blot lane loading control for (A). (B) Western blot analysis of cells transfected with ThBD(FLAG) plasmid with protein detection as described for (A). (C) ThBD/NRP2 DKO REPI cells transfected with plasmid guinea pig ThBD (red) or NRP2 (green) then infected 24 h later with GPCMV(PC+) at MOI 1 pfu/ml. Infection of wild type REPI cells served as a positive control (blue bar) and DKO REPI cells (black) as negative control . At 4 dpi, cells were harvested, and virus yield was titrated on GPL cells. Results plotted as mean viral titre±sd vs. cell type for each plasmid transfection. (D)-(F) Human receptors and GPCMV infection on ThBD/NRP2 DKO REPI cells. (D)-(E) Western blot of plasmid transfected cells for human receptors huNRP2 (D) and huThBD (E) on REPI cells with Western blot detection of FLAG tagged proteins as described for (A) and (B) above. (F) ThBD/NRP2 DKO REPI cells transfected with plasmid human huThBD (red) or huNRP2 (green) then infected 24 h later with GPCMV(PC+) with virus titer determined as described for (C) compared to infected control cells ThBD/NRP2 DKO (black) and REPI cells (blue). Statistical analysis Student *t* test, * *p* < 0.05; ns = non-significant.

### 7.3 Guinea pig CD46 is not a PC specific receptor and can bind GPCMV gH/gL complex

In addition to ThBD and NRP2, an additional receptor (CD46) has been identified for HCMV cell infection (46). However, HCMV studies are ambiguous regarding CD46 and PC-based cell entry since ectopic CD46 expression was unable to rescue HCMV infection on NRP2 knockout retinal epithelial cells unlike ectopic ThBD (47). Additionally, the inability of soluble CD46 to compete for PC binding further suggests that CD46 is not a PC specific receptor (47). The guinea pig encodes a CD46 homolog protein with 53% identity to human CD46 protein based on BLAST alignment. However, guinea pig CD46 expression has limited tissue expression and mainly found in the testis (59). In order to determine if guinea pig CD46 interacted with GPCMV glycoprotein complexes, transient plasmid CD46 (FLAG epitope tag) expression studies were carried out with GPCMV gH/gL, trimer (gH/gL/gO) or PC in separate IP assay studies using GFP-tagged gH as previously described (23, 52). Although CD46 was detected as an IP product in trimer and PC assays (Fig 5), it was also immunoprecipitated by gH/gL and demonstrated that guinea pig CD46 specificity was to the basic gH/gL complex (Fig 5B). In contrast, gH/gL had no interaction with endogenously expressed ThBD (Fig 5B). A control IP assay with GFP and CD46 failed to demonstrate any interaction with CD46 (Fig 5E). Results suggest that guinea pig CD46 might function as a co-receptor for infection via any gH/gL complex on the viral particle. To further determine the impact of CD46 on GPCMV infection, we evaluated comparative effect of CD46, NRP2 and ThBD ectopic expression on PDGFRA/NRP2 DKO GPL cells, which are resistant to GPCMV. Ectopic NRP2 or ThBD restored GPCMV infection to normal level of wild type GPL cells but CD46 failed to have any effect (Fig 6A & B). Additionally, ectopic expression of CD46 on ThBD/NRP2 DKO REPI cells did not enhance infection compared to ectopic NRP2 or ThBD which restored GPCMV infection levels to that of wild type REPI cells (Fig 6C & D). We concluded that ectopic over-expression of CD46 does not enhance GPCMV infection of fibroblasts or epithelial cells.

**Figure 5.**
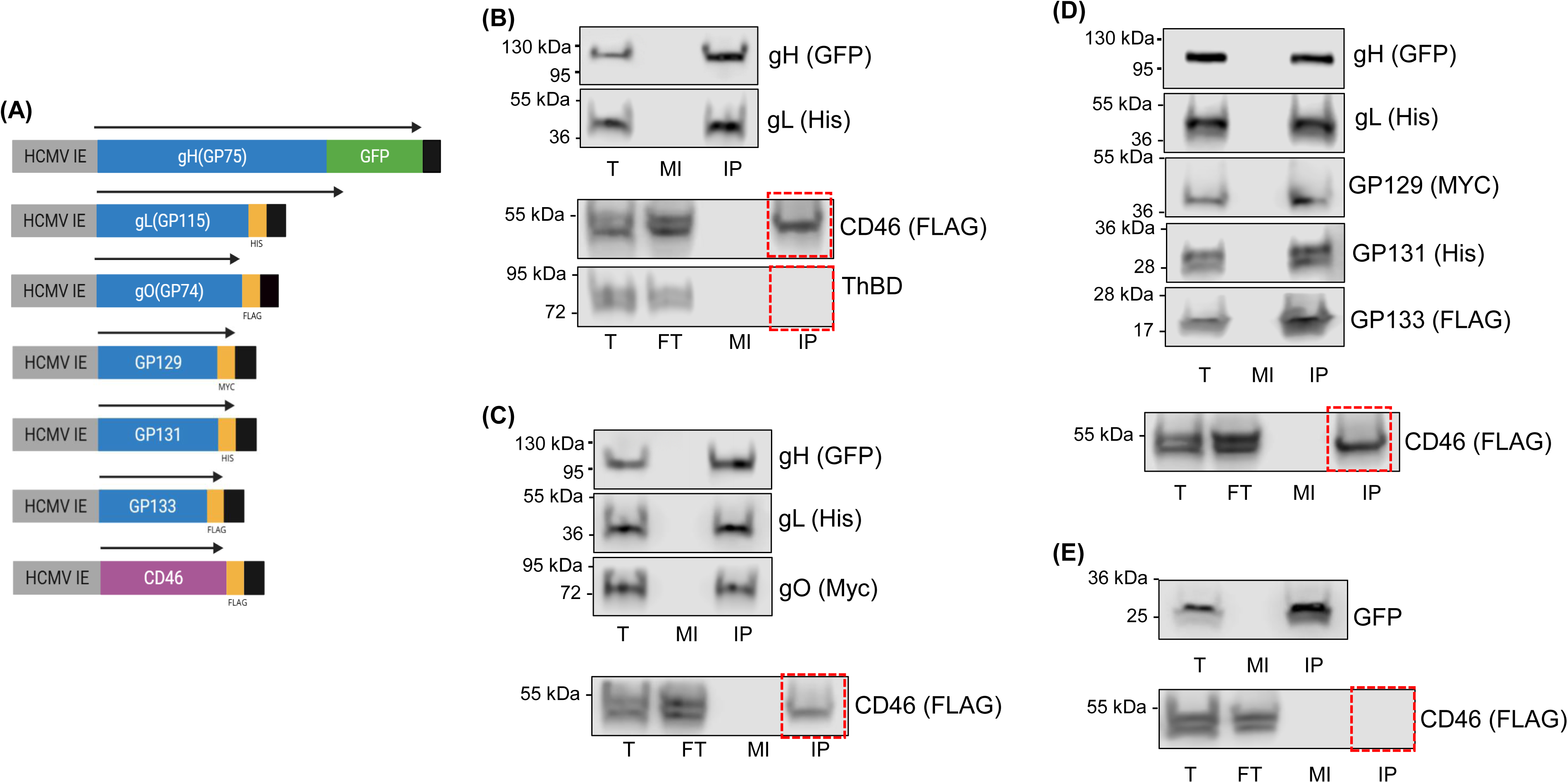
GPCMV gH-based trimer or pentamer complex or gH/gL and CD46 IP assay. (A) Diagram of specific expression plasmids, gene cassettes used for individual trimer and PC components and CD46. (B) Determination of CD46(FLAG) plasmid expressed protein interaction with gH/gL Receptor and complex expression and GFP trap IP as previously described (52). Primary antibodies used in Western blot analysis: α-GFP (gH), α-HIS (gL), α-FLAG (CD46), α-ThBD (ThBD). Determination of CD46(FLAG) plasmid expressed protein interaction with: (C) gH-based trimer; or (D) gH-based pentamer. Receptor and complex expression and GFP trap IP as previously described (52). Primary antibodies used in Western blot analysis: α-GFP (gH), α-HIS (gL), α-MYC (gO), α-MYC (GP129), α-HIS (GP131), α- FLAG (GP133), α-FLAG (CD46). (E) Control IP of GFP and CD46 co-expression to demonstrate lack of CD46 interaction with GFP. Samples were analyzed as total cell lysate or processed for IP assay, followed by Western blot analysis. T, total cell lysate; F, flow-through; MI, mock control lysate; IP, immunoprecipitated product.

**Figure 6.**
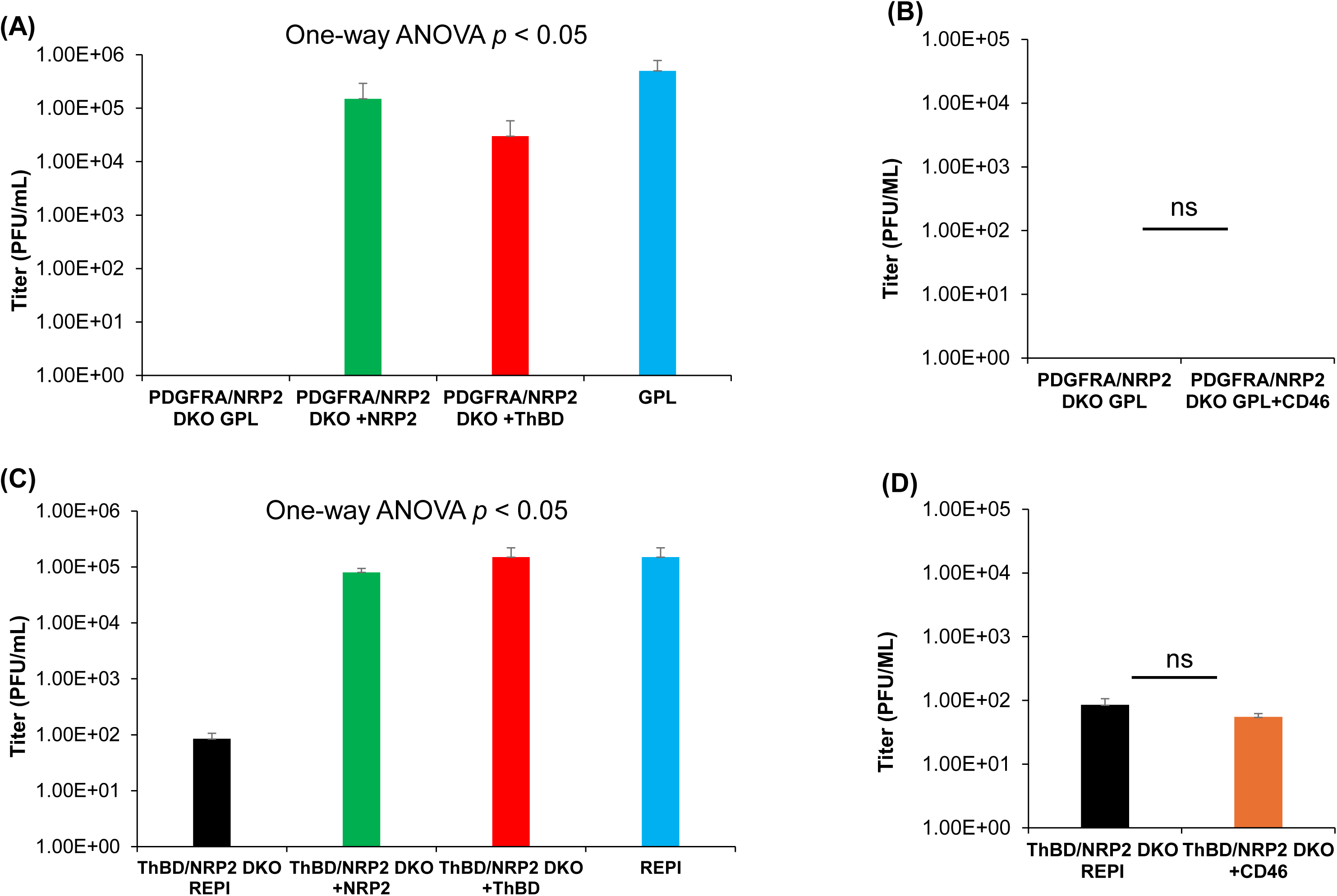
Ectopic ThBD complements GPCMV infection on receptor knockout fibroblast and epithelial cell lines unlike CD46. A-B) Fibroblast GPL cells. (A) GPCMV(PC+) growth (MOI 1 pfu/cell, 3 DPI ) on PDGFRA/NRP2 DKO GPL cells (black) compared to DKO cells ectopically expressing either NRP2 (green) or ThBD (red). Control wild type GPL(blue). One-way ANOVA *p* < 0.05. (B) GPCMV(PC+) growth on PDGFRA/NRP2 DKO GPL (black) compared to PDGFRA/NRP2 DKO GPL + ectopic expression of CD46 (orange). Both cell lines had minimal support for GPCMV infection. Student *t* test, ns = not significant). (C-D) Epithelial REPI cells. (D) GPCMV(PC+) growth (MOI 1 pfu/cell, 3DPI) on ThBD/NRP2 DKO REPI cells or DKO cells ectopically expressing NRP2 (green) or ThBD (red). Control wild type REPI (blue). One-way ANOVA *p* < 0.05. (D) GPCMV(PC+) growth on ThBD/NRP2 DKO GPL (black) compared to PDGFRA/NRP2 DKO GPL + ectopic expression of CD46 (orange). Student *t* test, ns = not significant).

### 7.4 Fibroblast derived GPCMV stock virus lacks full tropism to epithelial cells

Although cellular receptor expression may impact virus tropism, adequate surface expression on viral particles of GPCMV glycoprotein complexes associated with cell entry is also a factor. Mutant GPCMV(PC-) virus lacking one or all of the unique components of the PC (GP129, GP131 or GP133) (23) are highly impaired for infection of epithelial and endothelial cells as well as trophoblasts (23, 28, 49). However, wild type virus stock generated on GPL fibroblast cells is also potentially impaired for full spectrum of virus cell tropism. This is despite virus encoding PC when compared to GPCMV cell tropism of virus stock generated on epithelial cells. In order to illustrate this point, GPCMV(PC+) virus stock generated on renal epithelial (REPI) cells was compared to virus stock generated from a single passage on GPL fibroblast cells. GPCMV(PC+) GPL stock and GPCMV(PC+) REPI stock virus had similar growth kinetics on wild type GPL cells (Fig 7A). However, the GPL stock virus was highly impaired for infection of PDGFRA KO GPL cells unlike REPI virus stock (Fig 7A). This indicated a requirement for cellular PDGFRA expression for virus infection by GPL derived GPCMV stock virus (Fig 7A). Infection studies on epithelial cells demonstrated that for both renal epithelial and trophoblast cells, REPI derived virus stock had similar normal one-step growth curves with input virus MOI of 1pfu/cell (Fig 7B). In contrast, GPL derived virus stock was impaired for infection of both REPI and TEPI cell lines (Fig 7B). Overall, results demonstrate a significant difference in the ability of fibroblast or epithelial cell derived virus stocks to infect various cell types and dependence on PDGFRA receptor for cell infection. A single pass of GPCMV on fibroblast cells results in virus with impaired tropism for epithelial (Figure 7) and also endothelial cells (data not shown). Results indicate that the source of virus has an impact on virus cell tropism and potentially in virus pathogenicity in animal studies. Consequently, epithelial cell or animal derived salivary gland stock virus should preferentially be used for GPCMV animal research to maximize initial challenge virus cell tropism.

**Figure 7.**
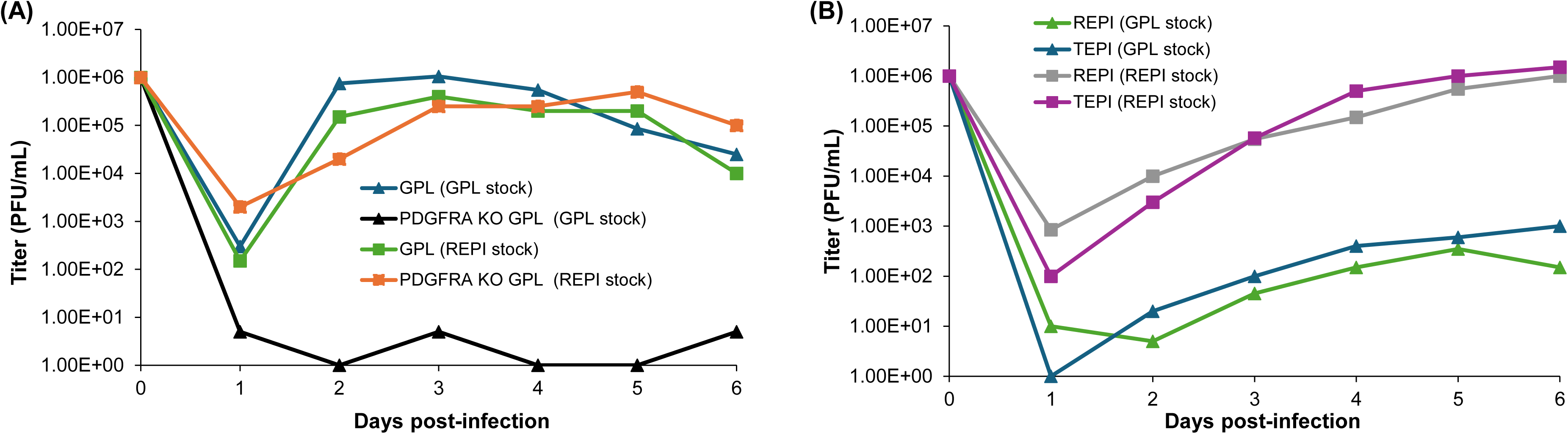
Cell type used (fibroblast v epithelial cells) for generation of virus stock impacts GPCMV cell tropism. Wild type GPCMV virus stock generated on REPI cells (REPI stock) or by single passage on fibroblast GPL cells (GPL stock) were evaluated for ability to infect GPL fibroblasts, REPI and trophoblast (TEPI) epithelial cells. (A) Comparative one-step growth curve of GPCMV (MOI 1 pfu/cell) on GPL and PDGFRA KO GPL cells. Viral titers evaluated for 1-6 days post infection (dpi). GPCMV GPL stock (triangle), GPL cell infection (blue), PDGFRA KO GPL cells (black). GPCMV REPI stock (square), GPL cell infection (green), PDGFRA KO GPL cells (green). (B) Comparative one-step growth curve of GPCMV on REPI or TEPI cells by virus stock generated on GPL cells (triangle) or REPI cells (square). Experimental set up similar to (A) with MOI of 1 pfu/cell. GPCMV (GPL stock) infection of REPI cells (green triangle), or TEPI cells (blue triangle). GPCMV (REPI stock) infection of REPI cells (grey square), or TEPI cells (purple square). Viral titers (1-6 dpi) were determined on GPL cells as described for A. Results shown are average values from replicates carried out in triplicate.

### 7.5. Epithelial cell receptor knockout does not enhance gB antibody neutralization

A gB vaccine is considered a corner stone of a cCMV vaccine because of the essential nature of gB and that it is an immunodominant neutralizing antibody target. However, the majority of gB antibodies against both HCMV and GPCMV gB are non-neutralizing but the use of a trimeric-gB rather than monomeric-gB vaccine antigen greatly enhances the neutralizing titer against GPCMV (33). Although this approach improves virus neutralization on fibroblast and epithelial cells, it does not completely block epithelial cell infection. We investigated if the knockout of PC receptor NRP2 or double-knockout of NRP2 and ThBD receptors had the potential to improve the neutralizing titer of historical gB vaccine sera from a prior study with trimeric capable AdgB vaccine (33). Additionally, we evaluated if knockout of the direct entry receptor (PDGFRA) on fibroblast cells (PDGFRA KO GPL) reduced neutralizing titer of gB vaccine sera as GPCMV infection could only occur by endocytic entry. Results in Figure 8 demonstrated that single or double-knockout of PC entry receptors on epithelial cells had no impact on improving gB vaccine sera neutralizing titer. Additionally, knockout of PDGFRA on GPL fibroblasts significantly reduced gB neutralizing antibody titer to level more similar to epithelial cells (Figure 8). A similar result was also obtained for Hartley GEFh v GEFh PDGFRA KO fibroblast cells (data not shown). Overall, it was concluded that the gB vaccine sera had reduced ability to neutralize GPCMV infection via endocytic pathway compared to direct entry. Additionally, the presence or absence of NRP2/ThBD endocytic pathway entry receptors did not impact gB antisera neutralization of GPCMV on epithelial cells. Possibly a limitation of the gB vaccine sera was a low-level antibody response to prefusion gB and increased response to this antigen may improve neutralizing titers (34) but awaits additional study.

**Figure 8.**
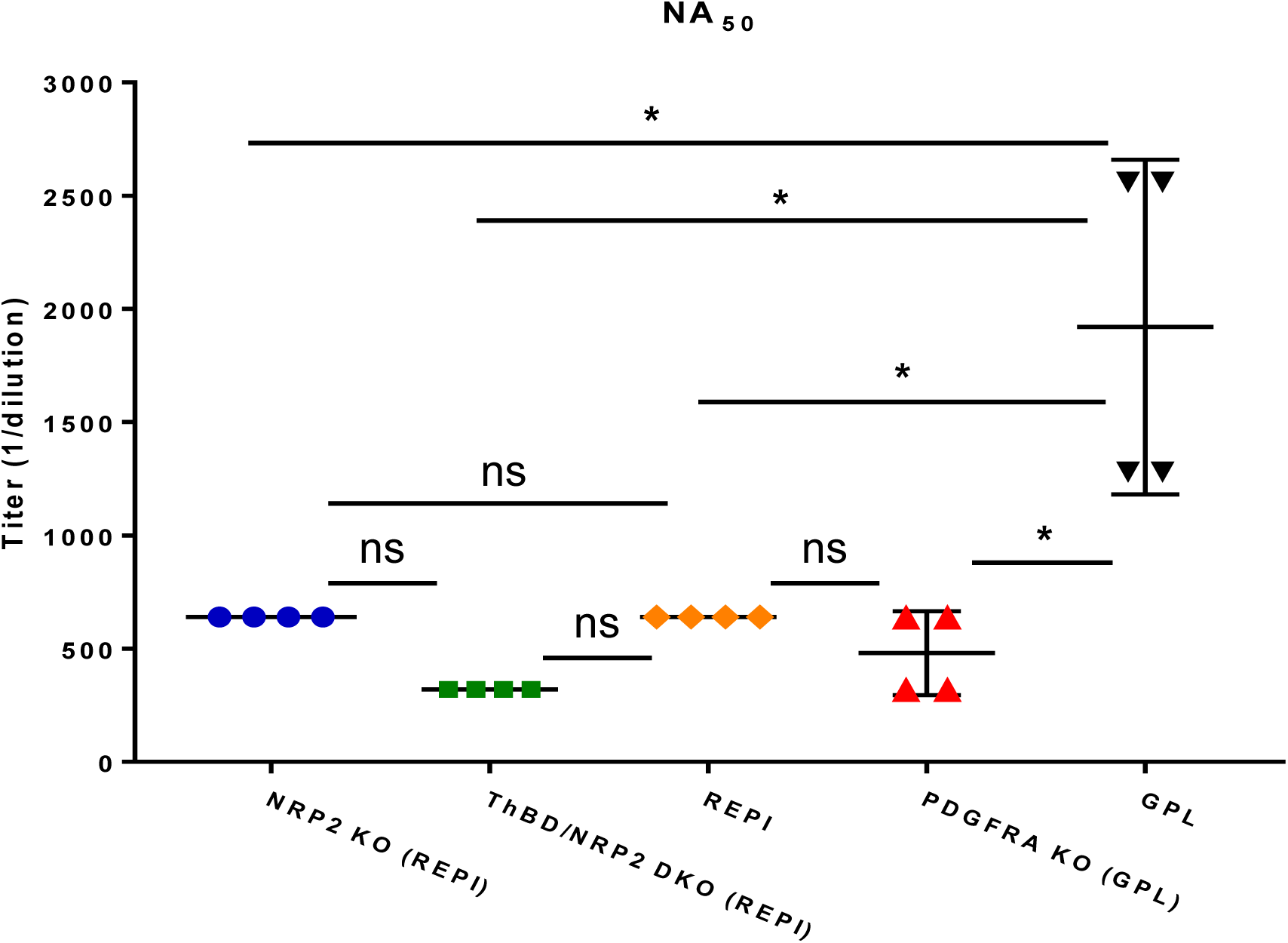
GPCMV neutralization (NA_50_) by anti-AdgB pooled animal sera on NRP2 KO and ThBD/NRP2 DKO compared to REPI cells. Pooled sera collected from animals inoculated with AdgBwt (n = 3) were evaluated for its ability to neutralize GPCMV PC+ wildtype virus on NRP2 KO REPI (blue circle); ThBD/NRP2 DKO REPI (green square); REPI (orange diamond) compared to PDGFRA KO GPL (red triangle) or GPL (black upside-down triangle). Student *t* test \**p* < 0.05; ns = non-significant.

## 9. Discussion

GPCMV PC is important for viral dissemination and pathogenicity in the guinea pig including congenital infection (23, 28, 49) and likely required to target monocytes and establishment of latency (50). HCMV, PC-based endocytic cell entry is only partially understood and poorly defined in animal cCMV models (guinea pig and rhesus macaques with RhCMV). It should be noted that murine CMV (MCMV) does not encode a PC and there is no cCMV model and MCMV utilizes NRP1 as a generalized method of cell entry more similar to Epstein-Barr Virus (60, 61). This study sought to evaluate candidate receptors for GPCMV cell entry in order to gain a better understanding of GPCMV infection and investigate the similarity between HCMV and GPCMV infection. Our prior studies identified the importance of cellular receptors PDGFRA and NRP2 in facilitating GPCMV infection via direct or endocytic pathways respectively and our current findings re-emphasize the importance of these receptors as well as ThBD. Previously we demonstrated that CD147 has an indirect method of action on GPCMV endocytic infection similar to HCMV (45, 52). Consequently, CD147 was not further evaluated in this current study since it did not directly interact with PC.

GPCMV trimer specifically interacts with PDGFRA in a species-specific manner but has no interaction with PC candidate receptors. Limitations of our previous GPCMV receptor study (52) were that experiments were performed on a fibroblast cell line to investigate both pathways of cell entry. Previously, Bafilomycin A treatment inhibited endocytic GPCMV entry in GPL PDGFRA knockout fibroblast cells, demonstrating a requirement for pH acidic flux similar to epithelial cells for GPCMV endocytic entry and requirement for NRP2 (52). In this current report, studies were expanded to epithelial cells to investigate endocytic infection on a cell type that required viral PC for successful infection. A further limitation of prior studies is that an ATCC cell line (GPL) of strain 13 guinea pig origin was utilized for GPCMV cell entry studies. This cell line was historically used for GPCMV infection studies by default as a convenient available embryo fibroblast cell line that easily supports GPCMV growth but is semi-transformed enabling extended passage of the cells unlike primary embryo fibroblasts. However, all GPCMV animal studies, including cCMV infections, are performed on Dunkin-Hartley strain animals with some limited studies performed on Strain 2 guinea pigs but not strain 13 animals (11). Furthermore, all established non-fibroblast guinea pig cell lines (both epithelial and endothelial) are derived from Dunkin-Hartley strain animals. Consequently, it was important to establish a Hartley derived fibroblast cell line and evaluate pathways of GPCMV cell entry. A new Hartley embryo fibroblast cell line (GEFh) was PDGFRA positive unlike all non-fibroblast Hartley cell lines established by our laboratory. PDGFRA knockout of GEFh cells blocked direct cell entry but not endocytic infection demonstrating both pathways exist in Hartley fibroblast cells.

In addition to the endocytic PC receptor NRP2, all Dunkin-Hartley cell lines (fibroblast and non-fibroblast) expressed a novel candidate receptor ThBD, which interacted with GPCMV PC but not gH/gL/gO trimer in immunoprecipitation assays. ThBD was absent from ATCC strain 13 GPL cell line and a lack of access to strain 13 animals prevented further study for tissue specific expression. Based on the Human Protein Atlas (https://www.proteinatlas.org/ENSG00000178726-THBD/tissue), ThBD is heavily expressed in endothelial cells but can be found on other cell types in humans and is present in high abundance in the placenta, including trophoblasts. Consequently, ThBD is likely an important receptor for transplacental HCMV infection. Presumably, based on ThBD function in endothelial cells, Kschonsak and colleagues (47) suggested that ThBD would likely be the HCMV PC receptor on endothelial cells and NRP2 the PC receptor for epithelial cells. However, their studies demonstrated ThBD virus receptor function on the backdrop of epithelial cells and exclusivity based on this cell type alone is unlikely if both receptors are expressed. Our analysis of human epithelial APRE19 cells detected both ThBD and NRP2 on these cells (data not shown). Guinea pig ThBD was expressed in embryonic fibroblasts, renal and amniotic sac epithelial cells, trophoblasts, umbilical cord vein and aorta endothelial cell lines, which suggests that ThBD along with NRP2 (52) are important receptors for virus dissemination and cCMV infection in Dunkin-Hartley guinea pigs. ThBD is more highly expressed in guinea pig endothelial cells than other cell types (Fig 2). Therefore, it is likely that ThBD is a key receptor for GPCMV endothelial cell infection. ThBD is also highly expressed in human neutrophils and likely an important receptor for HCMV PC dependent neutrophil infection with subsequent virus dissemination in the host but receptor expression in guinea pig neutrophils remains to be evaluated.

Although ThBD and NRP2 are important receptors for HCMV and GPCMV infection there is a species-specificity associated with these receptors that prevents human counterparts substituting as functional entry receptors in guinea pig cells. Presumably, the reciprocal situation is the case for HCMV infection despite conservation of functional domains. Ectopic expression of guinea pig receptors on ThBD/NRP2 DKO epithelial cell lines restored GPCMV infection to normal levels. Additionally, ThBD was functional as a receptor on PDGFRA/NRP2 DKO GPL fibroblast cells. Based on current and prior experiments, both NRP2 and ThBD are universally functional as GPCMV receptors in different cell types for endocytic cell entry and PDGFRA is similarly effective as a universal receptor for viral direct cell entry (26, 28, 52). Currently, it is unclear if NRP2 or ThBD is a more effective receptor for GPCMV PC as single-knockout of each receptor on epithelial cells resulted in similar infection kinetics for GPCMV compared to wild type REPI cells. Loss of both receptors had a more profound impact limiting GPCMV infection without any impact to control virus HSV-1.

Double-knockout of direct (PDGFRA) and endocytic (NRP2) receptors on GPL fibroblast cells completely blocked GPCMV infection (52). In contrast, DKO of endocytic receptors NRP2 and ThBD on epithelial cells did not completely block infection and there remained a low-level permissive infection on DKO NRP2/ThBD epithelial cells suggesting additional functional receptor(s) present on the cell. Previously, we had identified CD147 as an indirect or co-receptor for GPCMV PC endocytic cell entry (52). On GPL cells, single KO of CD147 had no impact on infection as GPL cells were positive for PDGFRA and direct entry was unaffected. GPL DKO of PDGFRA and CD147 blocked direct cell infection and also significantly impaired NRP2 dependent endocytic infection but this impact was not directly on surface expression of NRP2 (52). Since REPI cells and all established guinea pig epithelial cell lines are CD147 positive, it is likely that CD147 functions as an indirect GPCMV receptor to co-ordinate virion interaction of gB with EGFR or integrins but this awaits further study and limited by available antibodies for guinea pig specific studies. Although in HCMV, CD147 has been identified as an indirect receptor for HCMV endocytic infection the functional mechanism remains to be defined (45). However, CD147 is identified as a co-receptor for various viruses including alphaviruses, influenza and SARS-CoV-2 (62–64).

Although CD46 (also known as MCP) has been identified as an additional HCMV candidate PC (co-)receptor (46), immunoprecipitation studies with transiently expressed plasmid guinea pig CD46 with GPCMV viral glycoproteins demonstrated a specific interaction with gH/gL without a requirement for trimer or PC. Consequently, CD46, may function as a GPCMV co-receptor related to both pathways of infection but ectopic expression of CD46 on NRP2/ThBD DKO epithelial cells or NRP2/PDGFRA DKO fibroblasts had no impact on enhancing infection. It is possible that CD46 has limited function as a receptor in HCMV infection as studies by Kschonsak and colleagues (47) failed to demonstrate any HCMV infection complementation on NRP2 KO epithelial cells by ectopic CD46 expression whereas in contrast ectopic ThBD restored infection to wild type levels (47). Additionally, in HCMV studies, soluble CD46 had no impact on reducing epithelial infection unlike soluble NRP2 or ThBD (47). Currently it is unclear what level of gH/gL basic complex is present on the GPCMV virion surface compared to trimer and PC for CD46 interaction. However, since HCMV virions have gH/gL only complex in addition to trimer and PC, it is logical to assume that GPCMV similarly expresses gH/gL on virions. Since IP assays with gH/gL, trimer and PC were all successful in demonstrating interaction with CD46, potentially there is no requirement for gH/gL basic complex expression on GPCMV virions.

Based on current and prior research on GPCMV glycoprotein complexes and cell receptors in addition to current knowledge on HCMV and similarity to GPCMV, the overall strategy by GPCMV for infection via specific receptors for direct or endocytic cell entry pathways is based on various steps (A-D). (A) Regardless of entry pathway, GPCMV gM/gN glycoprotein complex is essential for initial virus cell attachment to heparan sulfate proteoglycans (HSPGs) on the cell surface prior to gH/gL based complex virus entry receptor engagement (22, 65). Knockout of either gM or gN is lethal to GPCMV demonstrating the essential nature of this complex (22). Subsequently, viral gH/gL based complexes in the form of the trimer (B) or PC (C) interact with respective receptors. This subsequently allows (D) gB modification from prefusion form to enable fusogenic gB based cell entry via EGFR interaction (24, 66). EGFR protein is expressed by all established guinea pig fibroblast, epithelial and endothelial cell lines (26) but confirmation of GPCMV gB/EGFR interaction awaits future study. Guinea pig and human EGFR share 89% identity based on BLAST alignment suggesting conservation of gB/EGFR interaction (26). It is unclear if cell signal transduction by EGFR or PDGFRA is required for gB mediated cell entry (67). In GPCMV, TK inhibitors had no impact on GPCMV infection of fibroblasts or epithelial cells (26). CD147 has an indirect undefined role to play in the endocytic viral entry pathway in both HCMV and GPCMV but KO did not modify surface expression of NRP2 (45, 52). Potentially, CD46 has a limited undefined role for CMV cell infection but this remains to be more fully investigated for both HCMV and GPCMV. Although gO is essential for trimer engagement of PDGFRA, knockout of gO (GP74) is not lethal to PC(+) virus where GPCMV retains the capability to spread by cell to cell contact without cell release virus and possibly in this infection strategy CD46 might have greater significance for gH/gL interaction (22, 23).

Possibly levels and ratios of GPCMV glycoprotein complexes might vary between cell types used for generation of viral stock. Most certainly, as demonstrated in the current study by infection, GPCMV PC levels on the virion differ between virus stock generated on fibroblast vs epithelial cells. Comparative studies of GPCMV (22122) viral stock generated (1x pass) on fibroblasts compared to GPCMV stock from renal epithelial cells demonstrated that fibroblast derived virus had restricted ability to infect epithelial, endothelial and trophoblast cells compared to epithelial derived viral stock (Figure 7). These results suggest that there is likely lower level of PC expressed on fibroblast derived progeny viral particles than epithelial derived virus stock. This is perhaps a cautionary warning against the use fibroblast generated GPCMV stock for pathogenicity or cCMV challenge studies. This would be extremely important for vaccine protection virus challenge studies where gB antibodies are disproportionally more effective against fibroblast infection compared to non-fibroblast cells (33) and more likely higher efficacy against fibroblast progeny virus. Original cCMV and pathogenicity studies were carried out with virus serially passaged in animals as salivary gland stock with this stock exhibiting similar tropism to epithelial cell derived tissue culture virus (11). However, fibroblast derived stock virus is likely to have reduced tropism to cell types in vivo as demonstrated by limited ability to infect epithelial and endothelial cells despite virus encoding PC. Potentially, challenge studies in animals with GPCMV fibroblast derived virus reduces translational impact of vaccine efficacy studies as only a percentage of virus would likely have an ability to infect non-fibroblast cells and fibroblast derived virus stock is more easily neutralized by gB antibody response.

In summary, this study extends GPCMV cellular infection studies with a focus towards endocytic infection and epithelial cell lines. In addition to NRP2, we verify that ThBD is a receptor for PC and endocytic cell entry with both receptors expressed on all established Dunkin-Hartley strain guinea pig cell lines including fibroblast, epithelial and endothelial cells. Both NRP2 and ThBD have the capacity to function independently as cell entry receptors with potentially expression levels favoring virus entry via one receptor compared to the other. In this regard, since endothelial cells express higher levels of ThBD compared to other cell types, this might be the main receptor for GPCMV endothelial cell infection. However, both NRP2 and ThBD are likely to be key receptors for placental infection given that trophoblast and endothelial cells express both receptors. Additionally, double-knockout of NRP2 and ThBD does not fully block GPCMV infection and suggests that additional receptors (eg. CD147) retain an indirect effect on endocytic pathway of infection but this awaits further study. Surprisingly, knockout of both NRP2 and ThBD on epithelial cells did not enhance neutralizing gB vaccine antibodies further demonstrating the importance of PC and a requirement for PC specific antibodies to block infection. Overall, results demonstrate similarity to HCMV and the continued importance of the GPCMV in the development of intervention strategies against cCMV in a high throughput model.

## 10. Author Statements

### 10.1 Author contributions

K. Yeon Choi: Conceptualization, Methodology, Investigation, Data curation, Formal analysis, Writing - original draft, Writing - review & editing. Yushu Qin: Investigation, Data curation, Formal analysis, Writing – original draft, Writing - review & editing. Alistair McGregor: Conceptualization, Methodology, Investigation, Data curation, Formal analysis, Writing - original draft, Writing - review & editing, Funding acquisition.

### 10.6 Acknowledgements

The authors would like to thank Alex Nguyen for his technical assistance in some tissue culture experiments.

---

**Figure S1. GPCMV and HSV-1 infection of guinea pig adenocarcinoma epithelial cell line GPC16.**

(A) Western blot analysis of ThBD and NRP2 expression on GPC16 cells compared to REPI and GPL and DKO cells. Upper blot (NRP2) cell lysate: GPC16 (lane1); REPI (lane 2), GPL (lane 3); ThBD/NRP2 DKO REPI (lane 4). Middle blot (ThBD) order as above. Lower blot gel loading confirmed by β-actin expression (order as above). (B) One-step growth curve of GPCMV(PC+) wildtype virus (MOI 1 pfu/cell) on GPC16 (blue circle) vs REPI (orange triangle) cells (1-6 days post infection). (C) HSV infection on GPC (blue) vs REPI (orange). MOI = 1, harvested 2 DPI. Statistical analysis Student *t* test \**p* < 0.05.

