## Supplementary material for "Cytomegalovirus pentamer dependent endocytic cellular infection utilizes redundant entry receptors in the guinea pig model": Suppl Fig1

Figure S1

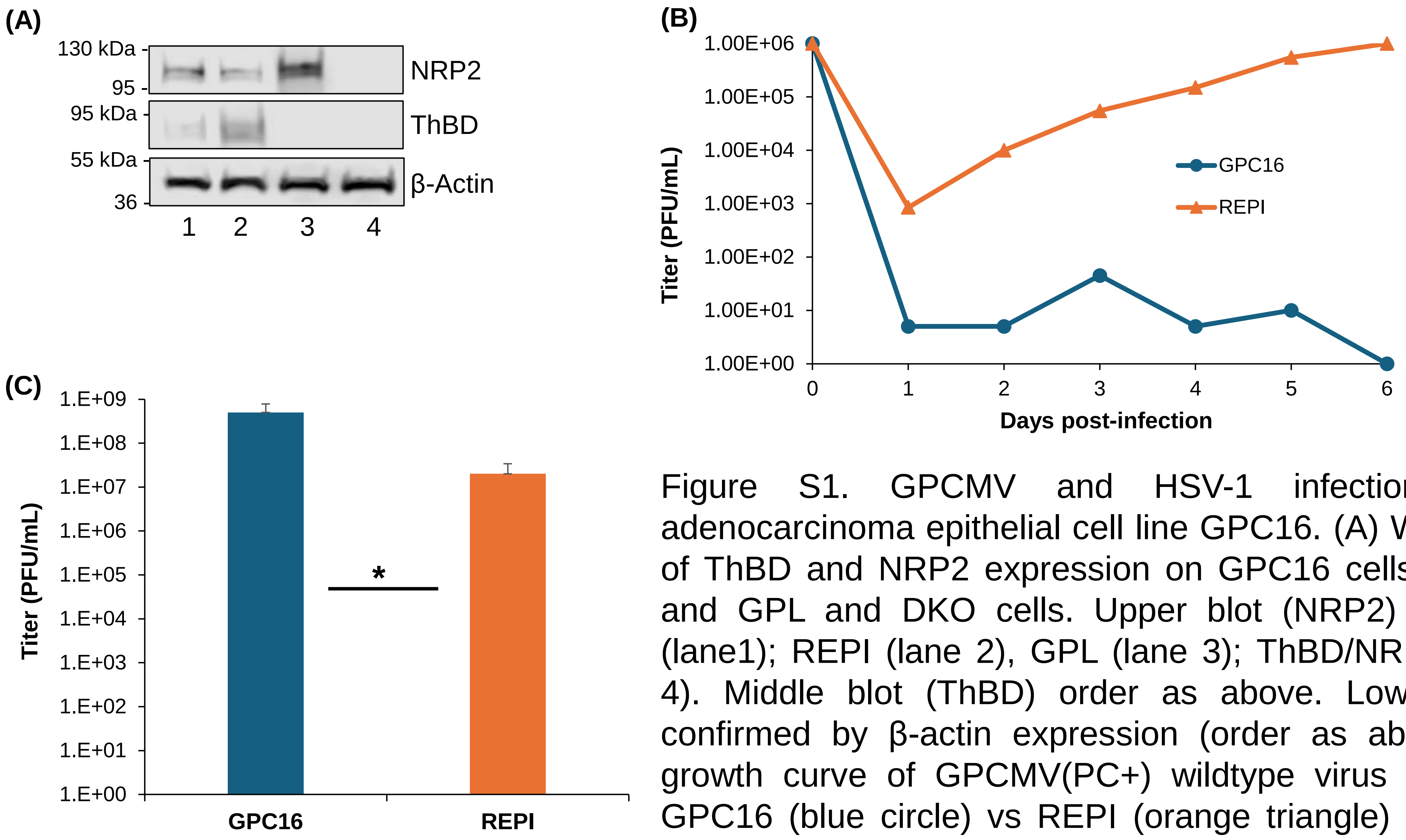


Figure S1. GPCMV and HSV-1 infection of guinea pig adenocarcinoma epithelial cell line GPC16. (A) Western blot analysis of ThBD and NRP2 expression on GPC16 cells compared to REPI and GPL and DKO cells. Upper blot (NRP2) cell lysate: GPC16 (lane1); REPI (lane 2), GPL (lane 3); ThBD/NRP2 DKO REPI (lane 4). Middle blot (ThBD) order as above. Lower blot gel loading confirmed by β-actin expression (order as above). (B) One step growth curve of GPCMV(PC+) wildtype virus (MOI 1 pfu/cell) on GPC16 (blue circle) vs REPI (orange triangle) cells (1-6 days post infection). (C) HSV infection on GPC (blue) vs REPI (orange). MOI = 1, harvested 2 DPI. Statistical analysis Student t test $p < 0.05$
